# Characterising ontogeny of quantity discrimination in zebrafish

**DOI:** 10.1101/2021.02.25.432846

**Authors:** Eva Sheardown, Jose Vicente Torres-Perez, Sofia Anagianni, Scott E. Fraser, Giorgio Vallortigara, Brian Butterworth, Maria Elena Miletto-Petrazzini, Caroline H. Brennan

## Abstract

A sense of non-symbolic numerical magnitudes is widespread in the animal kingdom and has been documented in adult zebrafish. Here we investigated the ontogeny of this ability using a group size preference task in juvenile zebrafish. Fish showed group size preference from 21 days post fertilization (dpf) and reliably chose the larger group when presented with discriminations of between 1 vs. 3, 2 vs. 5 and 2 vs. 3 conspecifics but not 2 vs. 4 conspecifics. When the ratio between the number of conspecifics in each group was maintained at 1:2, fish could discriminate between 1 vs. 2 individuals and 3 vs. 6, but again, not when given a choice between 2 vs. 4 individuals. These findings are in agreement with studies in other species suggesting the systems involved in quantity representation do not operate separately from other cognitive mechanisms. Rather they suggest quantity processing in fish may be the result of an interplay between attentional, cognitive and memory-related mechanisms as in humans and other animals. Our results emphasise the potential of the use of zebrafish to explore the genetic and neural processes underlying the ontogeny and function of number cognition.

## Introduction

One aspect of quantity discrimination is the evaluation of the number of items in a group, the numerosity. Although quantitative abilities of many species have been studied ^1–3^, research into the ontogeny of the ability to assess numerosity has been restricted so far to humans ^4,5^, fish (*Poecilia reticulata*), and domestic chicks (*Gallus gallus*) ^6,7^. Typically, in species other than chicks, which are a precocial species, the ability to assess numerosity changes with age until individuals reach adulthood. However, one day old guppies (*Poecilia reticulate*) can distinguish between a small number of conspecifics in a similar way to 40-day-old guppies ^8^.

Interestingly, guppies show no ratio effect for contrasts between numerosities 1 to 4, but the familiar ratio effect, Weber’s Law, for larger numerosities ^9^. Similar findings have been observed in human infants ^10–14^ suggesting the existence of conserved mechanisms of numerosity discrimination, though little is known about either the neural or genetic bases of these abilities (for a recent review see Lorenzi, E., *et al.* (2021)^15^).

The observation that small number discrimination does not show a ratio effect but larger numerosity discrimination does has been taken as evidence for two mechanisms for assessing number ^16,17^: an object-tracking system (OTS) or subitizing system with an upper limit of 4 items, and an approximate number system (ANS) for numerosities greater than 4 ^18–21^. However, this has been challenged in both humans and other species [e.g. Gallistel and Gelman (1992, 2000) ^9,22^ have proposed an accumulator mechanism, following Meck & Church (1983) ^23^, that operates across the numerical range]. Nonetheless, studies in both humans and animals showed that continuous physical variables that covary with number (e.g. overall space occupied, amount of motion, etc.), rather than numerical information per se, may play a prominent role in discrimination tasks ^24–28^. However, whatever the mechanism, either numerical or quantitative, it may interact with other cognitive systems, such that the effects of ratio in a discrimination task could be affected by, for example, working memory or attention ^29^.

Although zebrafish has rapidly become a well-established model in neuro-behavioural research ^30^, studies have focused on visuo-motor behaviour ^31^, decision-making mechanisms ^32^, and non-numerical discriminative abilities ^33^. The ontogeny of quantitiy discrimination in this species has not been studied. Here, as part of a larger project investigating the genetic bases and neural circuitry of numerical capacities ^34–36^, we examined the ontogeny of quantity discrimination using a group size preference assay ^37–39^. This procedure has been widely used to investigate numerical competence in several fish species (e.g. guppy, mosquitofish, stickleback, red-tail splitfin) ^7,40,41^. As previous studies ^42^ suggest adult zebrafish utilise an approximate number system to represent both small and large numerosities, we presented juvenile fish with a choice between sets of individuals within the small (<4) number range as well as contrasts that span both small and large numbers.

## Methods

### Subjects

All subjects and stimuli were juvenile zebrafish bred in the fish facility at Queen Mary University of London. The fish were Tubingen (TU) wild type laboratory strain ^43,44^ In total ~2035 fish were used, ~1300 as stimulus fish, 535 as test fish (see individual experiments for n’s). At the time of the experiments the fish were 21-33 days post fertilization (dpf), ages chosen on the basis of previous studies suggesting zebrafish show group size preference from 3 weeks post fertilization ^45,46^. Prior to experiments fish were kept in large groups (40-50 fish per tank) of unknown sex in a recirculating system under standard rearing conditions ^47^. During rearing, fish were kept at a constant temperature of 28 ± 2°C, with a 14:10 light: dark cycle and fed twice a day with paramecium and Gemma 75/150 (Skretting, USA). Each subject was used only once. After testing all fish were returned to the facility and maintained as breeding stock.

### Procedure

The group size preference (GSP) assay was performed using experimental apparatus that ensured equal spacing between stimuli fish as described in Gómez-Laplaza and Gerlai (2012, 2013) ^40,48^ (Fig. 1a, b, c). Fish were fed an hour before testing and all experiments were conducted between 10am and 13.00pm. Stimuli and test fish were matched for age and size. Fish behaviour was recorded using an overhead camera in the Daniovision larval tracking system (Noldus, UK). The video recordings were analysed using Ethovision software (Noldus, UK)

**Figure 1.**
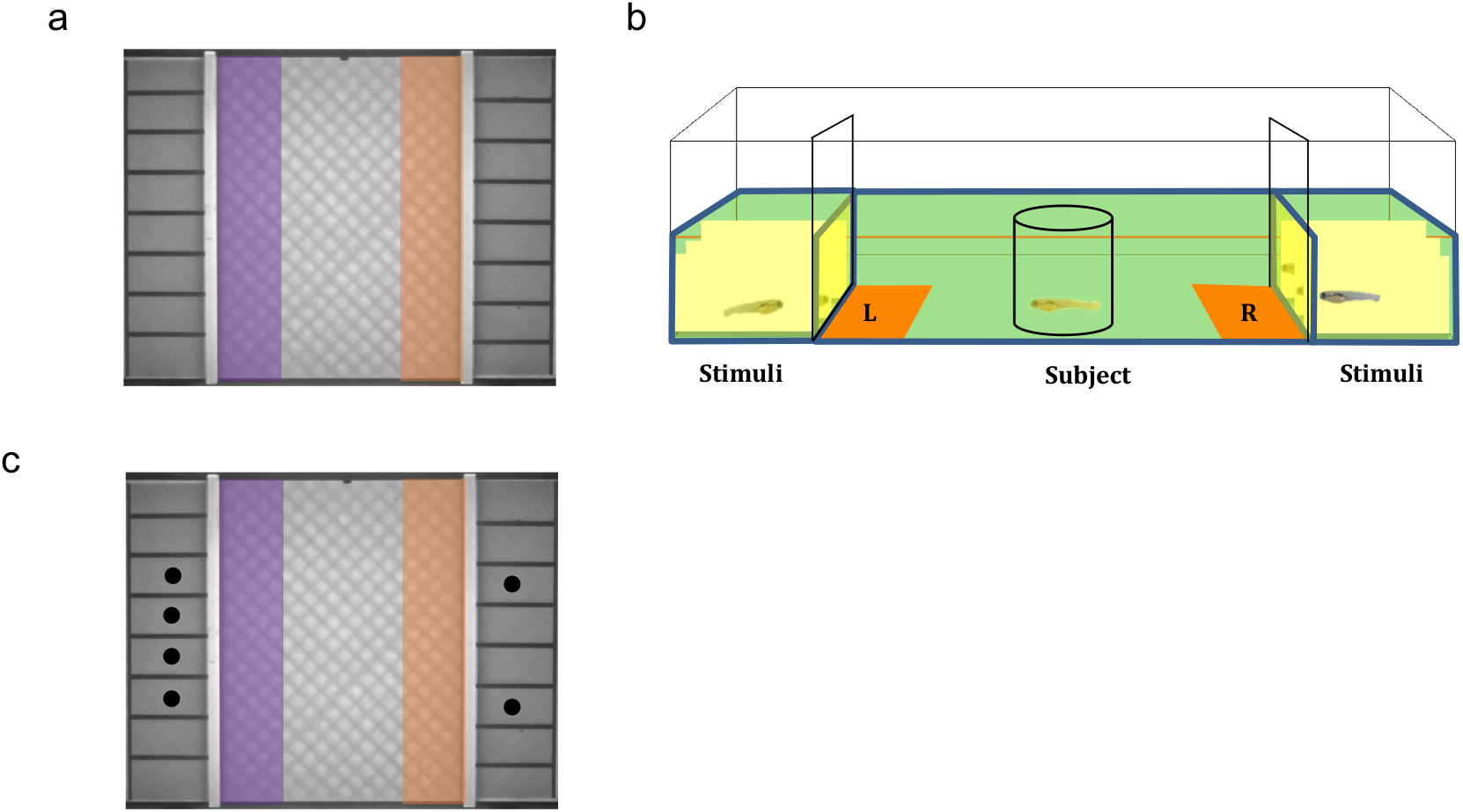
Screenshots (a, c) of apparatus and schematic representations of individual arenas (b). **a).** The set up consists of a single arena. Ethovision detects the presence of a subject in the left or right zones (indicated by coloured boxes), which sit 1cm in front of the stimulus windows. **b).** The arena is divided into three adjoining compartments. The central compartment is 8cm long and 6cm wide, with the 8cm sides being clear for viewing the stimulus compartments. Each stimulus compartment is divided into eight 1.5 cm long and 1cm wide opaque sections. At the beginning of the trial the subject is held in a removable transparent cylinder with a 2cm diameter. A removable opaque barrier was present between the stimulus fish and test fish. **c).** Schematic of ‘controlled space’ conditions with the white dots representing the positions of the outermost zebrafish in a controlled 2 vs. 4 contrast.

#### 1. Ontogeny of group size discrimination

As previous work reports robust social preference for ages of 21 dpf and above, we investigated the ontogeny of GSP zebrafish from 21-33 dpf. As slight variation in developmental rates make exact staging difficult, we pooled across days matching for size (21-23, 24-26, 27-29, 30-33 dpf). Stimulus fish were held in equal sized (1 x 1.5 x 1.5 cm) individual compartments as in Fig.1 a, c. Stimuli fish were introduced to their compartment with an opaque barrier between the stimulus fish and central compartment and allowed to habituate for 5 min. The test fish was introduced into the central compartment and allowed to explore the entire area for 5 min. After this time, the fish was restricted to a central domain using a clear plastic cylinder (diameter 2 cm) that was manually placed over the fish and used to usher it to the centre of the compartment (an etched circle indicated the centre of the central compartment). Once the test fish was in the centre, the barriers were lifted to reveal the stimulus fish. The test fish was allowed to view the stimulus fish for 1 min and then the holding cylinder was removed, and the behaviour of the test fish recorded over a 5 min period. We assessed performance in 1 vs. 3 (easiest), 2 vs. 5, 2 vs. 4, and 2 vs. 3 (most difficult) (N = 129 n= ~8 fish for each contrast and age group). The position of the larger group was counterbalanced across trials to control for any side bias. The output measure, the overall index, was calculated using the formula:

The overall index gives us a measure of the preference for the larger group.

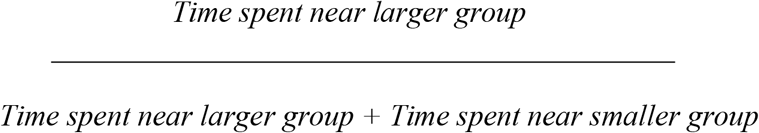

#### 2: Controlling for overall space

To test the hypothesis that fish were using the overall space occupied by the shoal as a discriminative cue rather than the number of conspecifics ^7,49^ we ran a control experiment where distance between the two most lateral fish was equal for both group sizes in 2 vs. 5, 2 vs. 4 and 2 vs. 3 discriminations. In this way, the outer stimuli fish on either side of the central compartment were positioned opposite each other (Fig 1c). We tested 30-33 dpf fish (N = 87, n=14-15 for each contrast) that were matched for age and size to stimulus fish.

#### 3: 1: 2 ratio discrimination when crossing the small and large number range

To test the hypothesis that proximity to the limits of the subitizing range (3 items) influences number discrimination, we assessed the ability of juvenile fish to discriminate a 1:2 ratio in the small number range (1 vs. 2) and when the numerical distance was increased from 1 to 3 (1 : (1 vs. 2), 2 : (2 vs. 4) or 3 : (3 vs. 6)) and the total number of items increased from 3 (1 vs. 2) to 9 (3 vs. 6). We included the contrast 2 vs. 5 as a positive control. We tested 30-33 dpf fish (N = 90 n = 20-25 for each contrast) that were matched for age and size to stimulus fish.

#### Statistics

All data significantly deviated from normality (Kolmogorov–Smirnov test, p < 0.05). We therefore used non-parametric statistics for all our data sets. Kruskal-Wallis tests followed by pair wise comparisons with Bonferroni correction were carried out to assess significant differences between groups. One sample Wilcoxon Signed Ranks tests statistics were used to compare preference for the larger group to chance level (0.50). All statistical tests were two tailed. Analyses were carried out in SPSS 27 and Graphpad Prism 9.

## Results

### 1. Ontogeny of group size discrimination

When we first analysed the ontogeny of the preference for the larger group across all contrasts we found no overall difference in preference for the larger shoal between different ages (χ^2^(3) = 2.835, *p* = 0.418). Analysis of difference in preference between contrasts with all ages found there was a significant preference between contrasts (χ^2^(3) = 10.060, *p* = 0.0158). Pairwise comparisons with Bonferroni correction, however, found no significant differences. One sample Wilcoxon signed rank tests showed that for the 1 vs 3 contrast there was a significant preference for the larger shoal across all age groups (21-23 dpf (*p* < 0.01); 24-26 dpf (*p* = 0.016); 27-29 dpf (*p* < 0.01); 30-33 dpf (*p* = 0.016)). This was also found with the 2 vs. 5 contrast, showing significant preference for the larger shoal across all age groups (21-23 dpf (*p* < 0.01); 24-26 dpf (*p* = 0.017); 27-29 dpf (*p* < 0.01); 30-33 dpf (*p* = 0.016)). There was no significant preference for the larger group in the 2 vs. 4 contrast at any age (p >0.05). There was no preference for 2 vs. 3 at 21-26 dpf (*p* >0.05), however there was significant preference at 27-29 dpf (*p* = 0.016) and 30-33 dpf (*p* = 0.016) (Fig. 2a).

**Figure 2.**
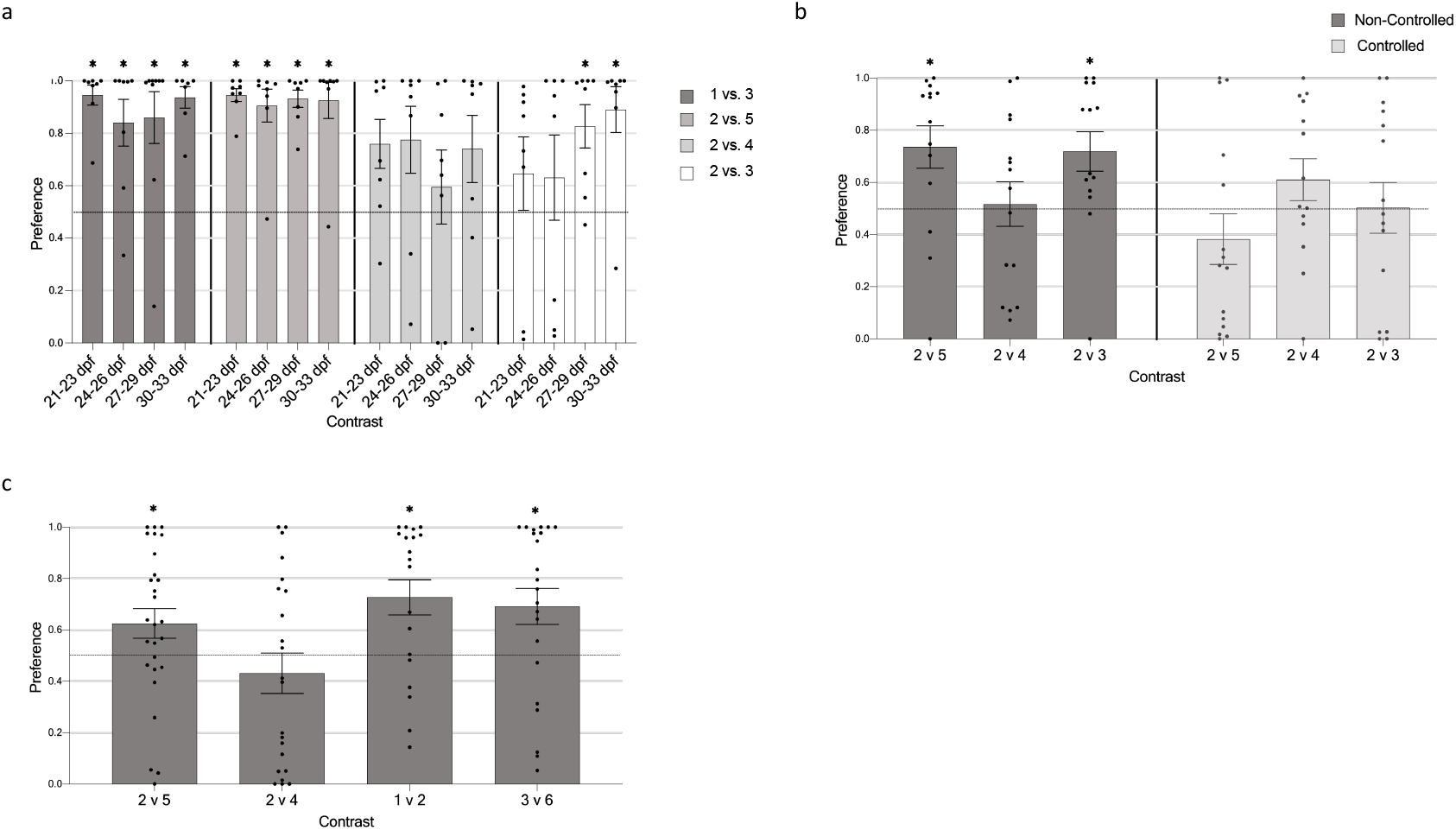
Investigating quantity discrimination. a). Ontogeny and limit of discrimination abilities. The fish of all ages showed discrimination of easier contrasts, 1 vs. 3 and 2 vs. 5 (respectively, 0.3 and 0.25 ratio), whereas a developmental change was observed when the most difficult comparison was presented: 2 vs. 3 (0.67 ratio). **b). Controlling for overall group space.** Without controlling for overall area (light grey columns), the fish show a significant preference for the larger group in both the 2 vs. 5 and 2 vs. 3 controls. Controlling for overall area results in absence of preference in all the contrasts **c). Investigating 1: 2 ratio.** Fish successfully discriminate 2 vs. 5, 1 vs. 2 and 3 vs. 6 but not 2 vs. 4. Dashed lines indicate chance level, asterisks indicate significant difference to chance with *p* <0.05.

### 2. Controlling overall space

When the overall space occupied by the shoal was controlled, a Kruskal-Wallis test (for ‘controlled space’ condition) showed there was a main effect of space condition (χ^2^(1) = 4.296, *p* = 0.038) such that when the overall space occupied by the shoal was controlled fish spent significantly less time with the larger group. One sample Wilcoxon signed rank tests showed a statistically significant preference for the larger group when compared to chance only in the non-controlled groups for 2 vs. 5 (*p* = 0.025) and 2 vs. 3 (*p* = 0.013) but not 2 vs. 4 (*p* >0.99) (Figure 2b).

### 3. 1: 2 ratio discrimination when crossing the small and large number range

A significant difference in preference between contrasts was found (χ^2^(3) = 8.631, *p* = 0.035) (1 vs. 2, 2 vs. 4, 2 vs. 5, 3 vs. 6). Although in the contrasts 1 vs. 2 and 3 vs. 6 fish appeared to show a greater preference for the larger group than in the 2 vs. 4 contrast, post hoc comparisons did not reveal a significant difference (1 vs. 2 and 2 vs. 4 (*p* = 0.07) and 3 vs. 6 and 2 vs. 4 (*p* = 0.07)). Juvenile fish showed a significant preference for the larger group in the 1 vs. 2 (*p* < 0.01), 3 vs. 6 (*p* = 0.011) and 2 vs. 5 (p = 0.025) but not 2 vs. 4 contrasts (Figure 2c).

## Discussion

Despite the adaptive value of quantitative abilities ^50^, research has focused on adults with developmental studies limited to a few species. Here we investigated quantitative skills of juvenile zebrafish using a ‘group size preference’ task in which the fish had to choose between two groups of conspecifics differing in number. Several fish species have been found to join the larger shoal when exploring a novel and potentially dangerous environment to reduce predation risk ^41^. However, despite adult zebrafish forming tight multimember groups, social preference only becomes robust at the third week ^51^.

When assessing the ontogeny of group size preference from 3 weeks of age, the fish of all ages spontaneously preferred the larger group of social companions at easier contrasts, 1 vs. 3 and 2 vs. 5 (respectively, 0.3 and 0.25 ratio), whereas a developmental change was observed when the most difficult comparison was presented: 2 vs. 3 (0.67 ratio). Indeed, zebrafish consistently selected 3 social companions over 2 only from the fourth week of life. These results are consistent with findings in humans, guppies, tadpoles and dogs where, although possessing some number sense at birth, numerical acuity increases across development ^7,8,52–54^. Hence, zebrafish possess some quantitative skills at an early age, but they are not mature enough to discriminate comparisons of increased difficulty. However, when a comparison of intermediate difficulty was used, 2 vs. 4 (0.5 ratio), none of the ages tested chose the larger shoal, which is surprising as fish ranging from 27 to 33 dpf were expected be able to select it based on their preference observed in the 2 vs. 3 contrast. Although at this stage of the study, no clear explanation could be provided for this result, the lack of preference observed at all ages could not simply be accounted for by a methodological flaw. For this reason, in the following experiments we included the 2 vs.4 contrast to further examine performance of fish in this contrast.

It is known that the numerosity of a set covaries with several continuous quantities, such as the amount of motion and the overall space occupied by the shoals, that can be used as a proxy to estimate which set is larger/smaller ^55^. Previous studies showed that fish can rely on non-numerical cues, including motion, to discriminate shoal size ^56,57^. In order to exclude this possibility, we used an established experimental set up that has been successfully used to study numerosity abilities in fish. The set up ensures that all stimulus fish are individually visible to the test fish, which is essential for optimal quantity discrimination ^40,48^: the stimulus fish were held in individual compartments that allowed little movement so that the subject fish could look at both groups from an equidistant position before making a choice. This setup not only let the subject have a global view of the two groups ^40,48^ but also limited fish movement and orientation, thus minimizing differences in stimulus fish activity. However, we cannot exclude the possibility that slight differences in overall motion may have affected quantity discrimination.

In relation to the covariable of overall space, when we controlled for this cue juvenile zebrafish did not choose the larger group of conspecifics. However, successful discrimination was observed in 2 vs. 3 and 2 vs. 5 but not in 2 vs. 4 when overall space was not controlled for. These results seem to suggest that the overall space occupied may have driven the choice for the larger group. Such a finding suggests that juvenile fish are unable to perform the discrimination based on numerical information alone. However, other non-numerical cues, such as overall size of the fish and degree of movement were not controlled for and may have influenced the results. Further, we cannot exclude the possibility that specifics of the apparatus, for example the visual separation of stimulus fish, influenced the outcome. Nonetheless, the fact that fish consistently failed to select the larger group when presented a choice between 2 and 4 individuals, whether controlled for space occupied or not, suggests the use of overall space is not the whole story, as 4 fish occupy twice the space. Further, even though other continuous variables may also contribute to fish numerosity discrimination (e.g. occupancy), use of continuous variables alone would not predict the failure of the 2 vs 4 contrast and success at 2 vs 3, as any other non-numerical variable should equally apply to all contrasts making the discrimination of 2 vs 4 easier than 2 vs 3. These data therefore support the hypothesis that zebrafish may use numerical as well as non-numerical information to represent number as in other species including humans ^24,58^. In respect to this issue, a theory of magnitude (ATOM) proposes there is a common magnitude sytem for space, time and number ^59^. This theory is supported by Webers’ law, which is seen in space, time and number across a range of species highlighting the similarity in cognitive processes for the three concepts with journals citing this previously ^60–62^. ATOM seems to suggest that if the zebrafish are using overall space as a quantitative cue, in theory they will be using the same system as for numerical discrimination therefore they can be using both modalities, space and number. ATOM, along with our evidence that the zebrafish perform 5 vs. 2 and 3 vs. 2 but fail with 4 vs. 2 confirms that they are not using only spatial cues, as if that was the case, they should successfully discriminate 4 vs. 2. Our results, where fish did not choose the larger group of 2 vs. 4 even when multiple visual cues were available, suggests there is some aspect of the underlying mechanisms that renders this specific comparison challenging. However, as we have not run extensive continuous controls, further experiments are needed to rule out other non numerical explanations.

Previous behavioural, neuroimaging and electrophysiological studies provided evidence of the existence of two distinct systems for processing numerosities in humans, the OTS and ANS ^16,29,63^. Some argue that the ANS alone is recruited for processing discriminations over the whole numerical range. Others, instead, argue that representations of small numbers (< 4) are carried out by the OTS whereas the ANS is activated solely for representation of large numbers (> 4) ^16,63^.

In animals, despite most of the studies suggesting that a single mechanism such as the ANS is involved in processing both small and large numbers ^62,64,65^, there is also evidence of two different systems ^7,42,66–69^. However, contrasting results in both humans and animals on the ability to discriminate quantities that crossed the small/large boundary (1 vs. 4) led to the suggestion that numerical processing, may be context-dependent ^9,10,14,49,70–75^.

In our study, when zebrafish were presented with three comparisons with the same ratio (0.5), they discriminated 1 vs. 2, 3 vs. 6 but not 2 vs. 4, thus violating Weber’s law, which states that magnitude discriminations are based on the ratio between the represented magnitudes ^22^. Hence, accuracy should decrease as the ratio increases, but not when the ratio is constant. Furthermore, discrimination did not follow the distance effect as successful discrimination was observed for a distance of 1 or 3 units but not 2. Finally, according to magnitude/distance effects, accuracy should decrease when the numerical distance is held constant but the numerical size of the contrasts increases, a condition we did not observe here as there was no difference in performance between 2 vs. 5 and 3 vs. 6. All together, these findings suggest that magnitude representation in juvenile zebrafish did not adhere to the principles of the ANS. However, juvenile zebrafishes’ ability to distinguish groups containing a large number of conspecifics (i.e. 6), suggests that they are equipped with a mechanism for processing them.

Cordes and Brannon (2009) ^75^ proposed two hypotheses to explain infants’ ability to discriminate small (< 4) versus large (> 4) numbers only when a sufficiently large ratio is given ^75^. According to the “noise hypothesis”, infants may first represent small sets through the OTS and large sets through the ANS and then convert object files into approximate magnitudes to compare the two quantities. In this scenario, the conversion from one representation to the other generates noise which increases with the number of items in the small set and necessitates a greater ratio for successful discrimination. Alternatively, according to the “threshold hypothesis”, infants may represent both small and large sets through the ANS, but below a ratio threshold, and within the small number range, the representations through the OTS trump the ANS because they are more precise and reliable. However, when the ratio between small and large numbers exceeds a threshold, small/large comparisons can be performed solely using the ANS. In this case, success should be consistently predicted by the same ratio and would not change as a function of the number of items in the small or large array. Neither of these hypotheses can explain our finding that juvenile fish can discriminate 2 vs. 5 and 3 vs. 6 but not 2 vs. 4 items as the noise hypothesis predicts that 3 vs. 6 creates more noise than 2 vs. 4 and hence a larger ratio would be required for successful discirmination, and the threshold hypothesis predicts that as 2 vs. 4 and 3 vs. 6 has the same ratio, there would be no difference in performance.

However, a third hypothesis suggests that the two systems are not differently specialized for small and large numbers but rather that attentional constraints and working memory determine if a set of items is represented as discrete individuals by the OTS or as an approximate numerical magnitude by the ANS ^29^. Here, the OTS would not be a mechanism specific for numerosity processing, rather it would be a more general system that allows parallel individuation of multiple objects that are stored in the working memory. Numerosity would then be indirectly inferred due to one-to-one correspondence between each real item and its mental representation ^21^. This mechanism seems to depend on attention as the number of items we can store in memory is related to the ability we have to attend to them as observed in studies showing that the capacity to precisely enumerate items is compromised in dual-task and attentional blink paradigms ^76–78^ suggesting that the ANS, that operates over the entire range of numerosity, is supplemented by an attention-based system, the OTS, specific for small number, that does not operate under demanding attentional load conditions ^29,77,79^. Our data are consistent with this latter hypothesis. The break-down of performance at 2 vs. 4 in all the experiments suggests that the maximum number of items juvenile zebrafish can attend and retain in memory is 5. When the groups exceed this limit, fish appear able to represent numbers as quantities through the ANS, provided that a minimum numerical distance is held, which seems to be 3 (2 vs. 5 and 3 vs. 6). However, when a contrast is close to the upper limit, we speculate that the fish may still try to focus on each item but then fail to keep track of them and be unable to activate the ANS as the difference between the two shoals is too small, thus leading to a failure to discriminate. Hence, here we suggest that the systems involved in quantity representation may not operate separately from other cognitive mechanisms, rather, that quantity processing may be the result of an interplay among attentional, cognitive and memory-related mechanisms that orchestrate numerical competence in zebrafish, as is hypothesised in both humans and other animals^29,80^. The findings that adult zebrafish can memorize and discriminate 2 vs. 4 conspecifics ^42^ indicate that the mechanisms underlying quantity representation change over development as is the case for humans. Future research investigating when this ability develops, and if changes in attentional capacity and working memory allow such comparison to be made effectively, can provide new insight into the complex mechanisms at the basis of quantitative and numerical competence.

Despite the zebrafish being a powerful model in the field of translational neuroscience research to investigate molecular mechanisms underlying neuropsychiatric disorders ^81–84^, a limited number of behavioural assays are available to investigate related cognitive deficits.. In humans, impairments of non-numerical abilities have been widely described in dyscalculia (a number and arithmetic learning disorder) ^17,85,86^ and evidence of familial aggregation suggests a genetic component in the evolution of this disorder ^87,88^. In recent years, the existence of an evolutionarily conserved non-verbal numerical system among vertebrates with a possible shared genetic basis has been suggested ^16,41^. In this context, the high degree of genetic homology with humans ^89^ makes zebrafish extremely useful to investigate the genetic mechanisms of numerical competence and their role in dyscalculia. Our study contributes to this aim as we provide a first behavioural tool for the assessment of quantitative abilities at an early age that can be employed for rapid screening of mutant lines for candidate genes potentially associated with poor numeracy. Further experiments finely controlling for continuous quantities will be essential to assess the extent to which zebrafish can purely rely on numerical information.

## Ethics

All animal procedures in this study were reviewed by the QMUL ethics committee (AWERB) and conducted in accordance with the Animals (Scientific Procedures) Act, 1986 and Home Office Licenses.

## Data availability statement

Supporting data files will be made public on dryad on acceptance. DOI: 10.5061/dryad.1vhhmgqs9.

## Author Contributions

CHB, MEMP, BB, SEF, GV conceived and designed the study. ES, MMP, JVPT, SA, RR conducted experiments. ES, MEMP, BB, GV, CHB analysed and interpreted data. All authors contributed to writing the manuscript.

## Acknowledgements

We thank Luca Galantini and members of the fish facility for technical support and Dr. Riva Riley for contributions to group discussions.

